# RelTime relaxes the strict molecular clock throughout the phylogeny

**DOI:** 10.1101/220194

**Authors:** Fabia U. Battistuzzi, Qiqing Tao, Lance Jones, Koichiro Tamura, Sudhir Kumar

**Affiliations:** Department of Biological Sciences, Oakland University, Rochester, Michigan; Center for Data Science and Big Data Analytics, Oakland University, Rochester, Michigan; Institute for Genomics and Evolutionary Medicine, Temple University, Philadelphia, Pennsylvania; Department of Biology, Temple University, Philadelphia, Pennsylvania; Department of Biological Sciences, Tokyo Metropolitan University, Hachioji, Tokyo, Japan; Research Center for Genomics and Bioinformatics, Tokyo Metropolitan University, Hachioji, Tokyo, Japan

## Abstract

The RelTime method estimates divergence times when evolutionary rates vary among lineages. Theoretical analyses show that RelTime relaxes the strict molecular clock throughout a molecular phylogeny, and it performs well in the analysis of empirical and computer simulated datasets in which evolutionary rates are variable. Lozano-Fernandez *et al.* (2017) found that the application of RelTime to one metazoan dataset (Erwin *et al.* 2011) produced equal rates for several ancient lineages, which led them to speculate that RelTime imposes a strict molecular clock for deep animal divergences. RelTime does not impose a strict molecular clock. The pattern observed by Lozano-Fernandez *et al.* (2017) was a result of the use of an option to assign the same rate to lineages in RelTime when the rates are not statistically significantly different. The median rate difference was 5% for many deep metazoan lineages for Erwin *et al.* (2011) dataset, so the rate equality was not rejected. In fact, RelTime analysis with and without the option to test rate differences produced very similar time estimates. We found that the Bayesian time estimates vary widely depending on the root priors assigned, and that the use of less restrictive priors produce Bayesian divergence times that are concordant with those from RelTime for Erwin *et al.* (2011) dataset. Therefore, it is prudent to discuss Bayesian estimates obtained under a range of priors in any discourse about molecular dating, including method comparisons.

## Introduction

RelTime was developed to estimate timetrees from molecular sequence data when evolutionary rates vary among lineages (Tamura *et al*. 2012, 2018). It has been shown to be accurate in the analysis of computer simulated data generated with extensive rate heterogeneity throughout the tree (Tamura et al. 2012; Filipski et al. 2014; Tamura et al. 2018). In analyses of many large empirical datasets, RelTime produced divergence times similar to those reported from Bayesian methods, when equivalent priors and calibrations were used (Mello et al. 2017). In addition, theoretical analyses clearly established that a relative rate framework, which does not assume a strict molecular clock, forms the mathematical foundation of the RelTime method (Tamura et al. 2018).

These theoretical and empirical findings are inconsistent with the Lozano-Fernandez *et al.* (2017) report, which concluded that RelTime functionally maintained a strict molecular clock for animal divergences in an analysis of one dataset containing 117 species and 2049 amino acids (Erwin *et al.* 2011). They surmised that this pattern is the cause of the curvilinear relationship between Bayesian and RelTime node age estimates observed by Battistuzzi *et al.* (2015). Lozano-Fernandez *et al.* (2017) also reported that the standard errors (SE) of RelTime estimates linearly increase in size with deeper node ages, unlike Bayesian approaches, and speculated that this pattern might explain the difference of animal divergence between Erwin *et al.* (2011) and Battistuzzi *et al.* (2015). Here, we present results from a re-analysis of Erwin *et al.* (2011) data to evaluate Lozano-Fernandez *et al.* (2017) concerns (**Fig. 1**). We estimated node ages and evolutionary rates using RelTime in MEGA (Tamura *et al.* 2013, Kumar *et al*. 2016) and Bayesian methods in Phylobayes (Pb) (Lartillot et al. 2009). These software were selected because they were used in the prior studies discussed here (Lozano-Fernandez *et al.* 2017; Battistuzzi *et al.* 2015; Erwin *et al.* 2011).

**Figure 1.**
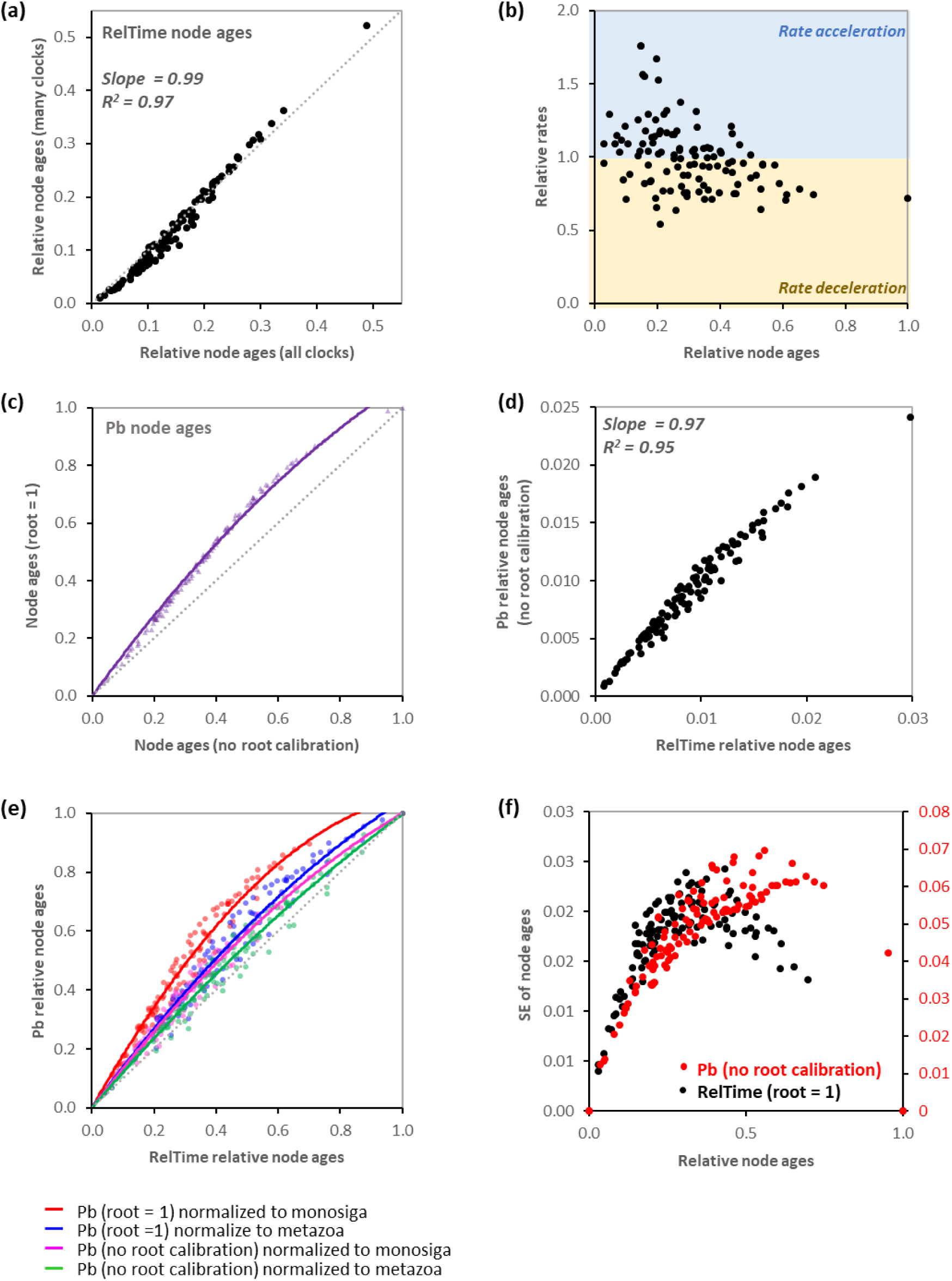
Comparisons of rates, dates, and standard errors from Bayesian and RelTme analyses. (**a**) RelTime estimates of node ages calculated with (“many clocks”) and without (“all clocks”) the rate merging option. The linear slope and *R*^2^ value are shown. Dotted gray line represents 1:1 relationship. (**b**) Normalized RelTime relative rates for nodes at different time depths, with rates greater than the average, above 1.0, showing acceleration and those below 1.0 showing a slow-down (blue and yellow backgrounds, respectively).Relative node ages were normalized to the age of ingroup root. (**c**) Relationship of Phylobayes node estimates without root calibration and with root age constraint at 1. Node ages were normalized to the age of the root. Solid line shows the polynomial fit and dotted gray line represents 1:1 relationship. (**d**) Relationship of RelTime and Phylobayes node ages obtained without root calibration. All node ages were normalized to the sum of ingroup node ages. The linear slope and *R*^2^ value are shown. (**e**) Relationship of RelTime estimates with Phylobayes with and without specified root calibration, and normalized to either Monosiga (Choanoflagellate) or Metazoa. Solid lines show polynomial fit for each comparison and dotted gray line represents 1:1 relationship. The R^2^ values for the polynomial fit are all greater than 0.94. (**f**) Standard errors (SEs) of node ages produced by RelTime and Phylobayes under different calibration constraints. Black circles: RelTime estimates of SEs of node ages when the ingroup root node is constrained at 1. Red circles: Phylobayes estimates of SEs of node ages without the root calibration; Phylobayes estimates were divided by 1000 for direct comparisons because root calibration is automatically set to 1000 when no root calibration is specified.

### RelTime relaxes the strict molecular clock in shallow as well as deep nodes

Lozano-Fernandez *et al.* (2017) stated that RelTime does not relax the strict molecular clock in the deep branches of a metazoan phylogeny, because the (relative) rates reported by RelTime were equal to 1 for many deep lineages; RelTime rates are all relative to the rate of the ingroup root node that is assigned a value of 1 for ease of reference. In RelTime, any deep or shallow lineage may receive the same rate as its ancestral lineage, when one chooses the “many clocks” option. Under this option, evolutionary rates of ancestral and descendant lineages are compared and merged if their equality cannot be rejected statistically (Tamura *et al.* 2012, 2018). This is indeed the case for the Erwin *et al.* data (see Figure 1c in Lozano-Fernandez *et al.* (2017)), where lineages near the root showed very similar evolutionary rates. However, this pattern is not unique to deep nodes. A few other lineages in shallower parts of the phylogeny also showed identical rates (e.g., 6 other lineages have the same relative rate of 1.86).

A RelTime analysis of another large dataset (274 species, 7370 sites; dos Reis *et al.* 2012), also using the “many clocks” option, confirms that rate identity is not influenced by the depth of the node, but rather the outcome of statistical tests carried out on all the nodes (**Fig. 2a**). In this dataset, lineages emanating from the ingroup root showed different rates, and many intermediate and shallow lineages showed similar rates (**Fig. 2a**). These rates were not statistically different, so they were assigned the same value. Importantly, in both of these datasets, the estimates of node ages showed an excellent linear relationship with and without the “many clocks” option (slope = 0.99 and 0.99 in **Figure 1a** and **2b**, respectively). A t-test did not reject the null hypothesis of equality of node ages (i.e., linear regression slope = 1) from RelTime analysis with many clocks and all clocks (*P*-value > 0.2). These results contradict Lozano-Fernandez *et al.*’s conclusion that RelTime imposed a strict molecular clock on deep divergences of the metazoan dataset, because the equality of rates they observed is the consequence of the lack of significant rate differences. Thus, RelTime does not impose a strict clock in deep divergences.

**Figure 2.**
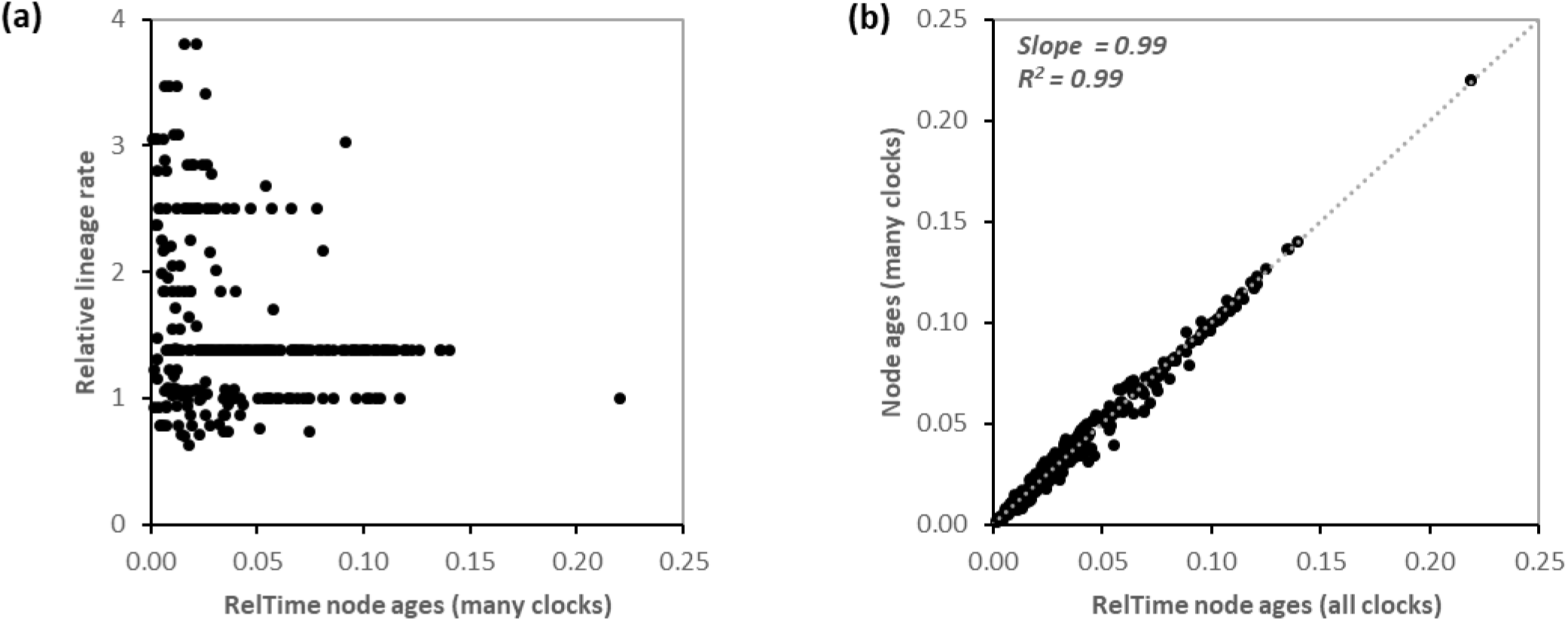
RelTime estimates of rates and relative node ages for a dataset of 274 species (dos Reis et al. 2012). (**a**) Rate estimates in relation to node ages obtained using “many clocks” option. (**b**) Comparison of node ages obtained with and without “many clocks” option. Slope and R^2^ values are shown.

Lozano-Fernandez *et al.* (2017) also commented on the rates produced by Bayesian and RelTime methods (see **Fig. 1c** and **1d** in Lozano-Fernandez *et al.* (2017)). These rates should not be compared, because Bayesian analyses produce evolutionary rates for individual branches in the phylogeny, whereas RelTime produces evolutionary rates for lineages (Tamura *et al.* 2018). A lineage rate is a function of all the branch rates in the subtree originating at the node of interest. Therefore, a lineage rate at any node in the tree is not expected to be equal to the evolutionary rate of specific branches directly connected to that node. In the case of Erwin *et al*.’s data, rates for many deep lineages were very similar, which may happen when branch rates increase and decrease over time within lineages. Their averages turn out to be very similar for some datasets (e.g., Erwin *et al.* 2011), but not others (e.g., dos Reis *et al.* 2012). We found that for nodes that were assigned the same rate of 1 in deep divergence, the median difference between ancestor and descendant rates was 5%.

In retrospect, detailed discussion of the similarity of time estimates obtained with and without the “many clocks” option in Battistuzzi *et al*. (2015) may have avoided the perception that RelTime is unable to relax the molecular clock for deep evolutionary timescales in the metazoan dataset (Lozano-Fernandez *et al.* 2017). The distribution of lineage rates obtained without using the “many clocks” option includes both higher and lower lineage rates throughout the tree (**Figure 1b**). Therefore, RelTime relaxes the strict molecular clock in deep as well as shallow lineages. Because RelTime node ages estimated with and without “many clocks” were very similar (**Figure 1a**), the relationship observed by Battistuzzi *et al.* (2015) between RelTime and Bayesian node ages cannot be caused by a lack of molecular clock relaxation. We, therefore, explored the possibility that the contrasting relationships between Bayesian and RelTime results observed by Battistuzzi *et al.* (2015) and Lozano-Fernandez *et al*. (2017) may be caused by the selection of priors in Bayesian analyses.

### Bayesian estimates with minimal sets of priors are not consistent with each other

Lozano-Fernandez *et al.* (2017) reported that RelTime estimates are “not proportional” to the Bayesian estimates obtained when they assigned an arbitrary root age calibration of 1 in their Phylobayes analysis with no clock calibrations. This analysis was meant to compare Bayesian and RelTime results using the minimum set of priors, because RelTime does not require specification of any priors or calibrations and produces relative times. Since the root age calibration of 1 is an arbitrary choice, we compared Bayesian estimates under an alternate minimum set of priors where no root age calibration was used (allowing it to default to 1000 in Pb, see **Methods** for details). The resulting relative node ages showed a curvilinear relationship between two sets of Bayesian estimates, with a 30% overall difference (**Figure 1c**). However, a Bayesian analysis using no root calibration and with a birth-death default prior produced time estimates that had a nearly linear relationship (slope = 0.97) with those from RelTime (Battistuzzi *et al.* 2015; **Figure 1d**). The similarity of results obtained between RelTime and Bayesian under a specific set of priors (i.e., birth-death and no root calibration) highlights the importance of discussing Bayesian estimates obtained under a range of priors in any discourse about molecular dating, including method comparisons, because Bayesian estimates can vary substantially depending on the prior choices (dos Reis et al. 2016; Inoue et al. 2010; Parham et al. 2012; Warnock et al. 2012, 2015, 2017; Hug & Roger 2007)

### Deep nodes are strongly affected by prior choices

As RelTime produces only relative time estimates, Bayesian time estimates must be scaled to a reference node for comparison with RelTime estimates. During our investigation above, we found that selection of the reference node used to scale times also influences the relationship observed between RelTime and Bayesian estimates. For example, the curvilinear trend obtained by using the age of the choanoflagellate Monosiga to normalize all ages for comparison (**Figure 1e**, red line) is very similar to that reported by Lozano-Fernandez *et al.* (2017) using the root node, but this trend becomes much less pronounced when Bayesian ages are normalized to the age of Metazoa, while maintaining a root calibration of 1.0 (**Figure 1e**, blue line). The relationship becomes even more linear when the root calibration is not specified (**Figure 1e**, pink and green lines). These results suggest that nodes are affected differently by prior selections and trends observed after normalization will be affected, depending on the node chosen for the normalization itself. In cases such as this one, when one or a few nodes have a strong impact on the identified trends, it may be advisable to normalize to the sum of time estimates of all nodes to allow every node to contribute to the normalization (e.g., Tamura et al. 2012).

### Bayesian and RelTime estimates of standard errors show similar trends

We also investigated whether the priors selected by Lozano-Fernandez *et al.* (2017) explain the reported differences between RelTime and Phylobayes standard error (SE) estimates for node ages in the metazoan dataset. While credibility/confidence intervals would usually be compared, here we present SEs to enable a direct comparison with values presented by Lozano-Fernandez *et al.* (2017). To generate results under comparable conditions, we fixed the ingroup root calibration to be 1 in RelTime. As expected, the relationship between node ages and SEs for RelTime showed a trend in which SEs first rise and then decrease with the increase of node ages (**Figure 1f**, black circles). This pattern is similar to that produced in the Bayesian analysis (see Figure 1f in Lozano-Fernandez *et al.* (2017)). This shows that the imposition of a root node calibration of 1.0 strongly affected SE estimates, because confidence intervals are clipped to avoid predating the calibration constraint. Upon omitting the root node calibration, we found that the Bayesian SE estimates increased with time (**Figure 1f**, red circles), similar to the pattern observed for RelTime without any constraints. Therefore, the decrease in SE with node age for Bayesian results observed by Lozano-Fernandez *et al.* (2017) was strongly influenced by the root node constraint, and was not directly comparable with RelTime results.

### Priors and calibrations that impact absolute molecular dates

Battistuzzi *et al.* (2015) used data from Erwin *et al.* (2011) as an example, because its analysis clearly showed that (a) the two maxima and the root prior have a very large impact on molecular time estimates produced by Bayesian methods and (b) different combinations of (maximum) calibrations and priors produce very different time estimates. Results similar to Battistuzzi *et al.* (2015) were also obtained by Lozano-Fernandez *et al.* (2017) (see their Fig. 3), but the induced effective prior was judged to be overly diffused.

These results highlight a well-known attribute of Bayesian analyses: the user-specified parameters and priors can be very different from the induced prior distributions due to complicated parameter interdependencies and relatively arbitrary approaches to truncation of the prior distributions (Warnock et al. 2015; Barba-Montoya et al. 2017; Eme et al. 2014). In the dataset analyzed here, it is clear that the Bayesian relative and absolute time estimates are strongly affected by the selection of priors. In the absolute dating analysis, the root prior used by both Erwin *et al.* (2015) and Lozano-Fernandez *et al.* (2017) is stricter than that explored in Battistuzzi *et al.* (2015). While the best prior cannot be unequivocally identified, a recent study on molecular timing of eukaryotes adds some new information. This study obtained a Bayesian divergence time estimate of ∼1375 million years ago for Opisthokonta (Animals+Fungi) in an analysis of a dataset with 116 taxa and 2166 amino acids (Gold et al. 2017). Using a dataset of comparable size, this study produced a divergence time estimate at the upper end of the marginal prior distribution used by Erwin *et al.* (2011) and Lozano-Fernandez *et al.* (2017), but well within the distribution of Battistuzzi *et al.* (2015) (see Table 2 in Lozano-Fernandez (2017)). Given the discordance among Bayesian analyses with different priors and RelTime, it is clear that the reliable establishment of the age of animal origin remains challenging.

## Conclusions

Lozano-Fernandez *et al.* (2017) made a specific conclusion about the ability of RelTime to estimate the timeline of animal diversification, which was based on an analysis of only one dataset. We have demonstrated that RelTime does not collapse rates, unless one chooses to assign equal rates to lineages that do not show statistically significant rate difference. Thus, RelTime is sensitive to rate changes in both deep and shallow portions of a phylogeny, which is consistent with the theoretical underpinnings of the RelTime method (Tamura *et al.* 2018). In fact, RelTime may be used as a reference framework to evaluate the effect of prior choices in Bayesian analyses. A RelTime analysis may also provide useful information about selection of priors and distributions used to diminish undue impact on molecular divergence times inferred using Bayesian methods, because, under some conditions, RelTime and Bayesian methods produced similar results for the data analyzed in Lozano-Fernandez *et al.* (2017). This complementary analysis may prove particularly useful in informing selection of priors for Bayesian analyses. Because Bayesian methods can become very computationally demanding, and RelTime speed is, in orders of magnitude, faster (Tamura *et al.* 2012, 2018), RelTime may serve as a practical and theoretically sound alternative to Bayesian methods for many datasets (Tamura *et al.* 2018).

## Material and Methods

The dataset consists of 117 species and 2049 aligned amino acids (Erwin et al. 2011). All analyses were conducted with RelTime (Tamura *et al.* 2012, 2018) and Phylobayes v. 4.1f (Lartillot et al. 2009). Phylobayes was selected because it was the Bayesian dating software used in all the previous studies discussed here (Erwin et al. 2011; Battistuzzi et al. 2015; Lozano-Fernandez et al. 2017). We also conducted RelTime analysis on another datasets consisting of 274 species and 7370 sites (dos Reis *et al.* 2012). For the RelTime analyses, no calibration times were used. All RelTime analyses were conducted with MEGA 6 (Tamura *et al.* 2013), the software used by Battistuzzi *et al.* (2015), with the exception of the SE calculation. The SE calculation implemented in MEGA 6 does not consider calibrations. Thus, to make the direct comparison with SE estimates in Pb analysis with a root constraint at 1 (**Figure 1f**), we conducted RelTime analysis in MEGA 7 (Kumar *et al.* 2016), which can accommodate user-specified calibration constraints. RelTime relative rates were normalized to their mean rate estimate over the whole tree for comparative purposes (**Figure 1b**). Note that RelTime produces relative lineage rates and these should not be directly compared to branch rates produced by Phylobayes. In the comparison of relative node ages obtained from RelTime with “many clocks” and “all clocks” options, we also conducted a t-test to examine whether the node ages from two options are significantly different from each other. i.e., the null hypothesis of regression slope is equal to 1 (*P* > 0.2). Two sets of parameters were used in Phylobayes analyses: one without any root calibration and another one calibrating the root node to 1 (as done in Lozano-Fernandez *et al.* (2017)). Phylobayes automatically scales node ages to 1000 when no root calibration is specified. We selected a birth-death speciation model with default parameters in these analyses. The final option used in this study was –cat –gtr –cir –bd with default hyperprior (10^−3^). All analyses in Phylobayes were run for at least 20,000 generations. While full convergence is expected to take many months, time estimates from these truncated analyses appear reliable, because relative time estimates remained stable at 2500, 6000, 8500, 12500, and 20000 generations. This convergence approach produced times identical to those obtained by Lozano-Fernandez *et al.* (2017) under the same analysis conditions and validated using Tracecomp (Lozano-Fernandez *et al.* 2017).

## Acknowledgements

We thank Heather Rowe and Beatriz Mello for comments and Davide Pisani for sharing the results from Lozano-Fernandez *et al.* (2017). We thank Nicolas Lartillot for guidance about the best ways to evaluate parameters in Phylobayes. We also wish to thank Davide Pisani for useful discussions and critical comments on the previous version of this manuscript, which does not imply his agreement with our conclusions. This work was supported by grants from National Aeronautics and Space Administration [NNX16AJ30G to F.U.B.], National Science Foundation [DBI 1356548 to S.K.], and Tokyo Metropolitan University [DB105 to KT].

